# Allele-specific RNA imaging shows that allelic imbalances can arise in tissues through transcriptional bursting

**DOI:** 10.1101/386359

**Authors:** Orsolya Symmons, Marcello Chang, Ian A. Mellis, Jennifer M. Kalish, Marisa S. Bartolomei, Arjun Raj

## Abstract

Extensive cell-to-cell variation exists even among putatively identical cells, and there is great interest in understanding how the properties of transcription relate to this heterogeneity. Differential expression from the two gene copies in diploid cells could potentially contribute, yet our ability to measure from which gene copy individual RNAs originated remains limited, particularly in the context of tissues. Here, we demonstrate quantitative, single molecule allele-specific RNA FISH adapted for use on tissue sections, allowing us to determine the chromosome of origin of individual RNA molecules in formaldehyde-fixed tissues. We used this method to visualize the allele-specific expression of *Xist* and multiple autosomal genes in mouse kidney. By combining these data with mathematical modeling, we evaluated models for allele-specific heterogeneity, in particular demonstrating that apparent expression from only one of the alleles in single cells can arise as a consequence of low-level mRNA abundance and transcriptional bursting.

## Introduction

Gene expression in genetically identical individual cells often deviates from that of the cell population average (Symmons and Raj 2016), which in mammals can impact cell fate and development (Abranches et al. 2014; Trapnell et al. 2014; Olsson et al. 2016; Mohammed et al. 2017), response to environmental stimuli (Cohen et al. 2008; Spencer et al. 2009; Halpern et al. 2017; Fritzsch et al. 2018) and disease (Shaffer et al. 2017; Tirosh et al. 2016; Avraham et al. 2015; Zanini et al. 2018). Over the past few years, it has emerged that at least some of this variability arises due to random fluctuations in the biochemical processes that underlie transcription and translation. In the case of transcription, a primary source of fluctuations is so-called transcriptional bursting, where a gene alternates between an active state, during which RNA is produced, and an inactive state, where no RNA is transcribed. Because both the time of onset of these bursts and the amount of RNA produced in a single burst are random, this process can lead to cell-to-cell variability (Raj and van Oudenaarden 2008; Vera et al. 2016; Nicolas, Phillips, and Naef 2017).

An additional nuance to the effects of bursting on cellular variability is that diploid mammalian cells carry two sets of chromosomes (one from each parent), which means that they also have two copies of each individual gene. It is typically assumed that for most genes both copies, called alleles, are capable of being expressed, thus providing protection through redundancy if one of them is mutated (Perrot, Richerd, and Valéro 1991; Sagi and Benvenisty 2017). Recent studies, however, which made use of crosses between distantly related mouse strains and high-throughput sequencing, uncovered that there can be extensive differences in the relative expression levels of the two alleles (Tang et al. 2011; Goncalves et al. 2012; Borel et al. 2015; Bonthuis et al. 2015; Andergassen et al. 2017). Additional work then showed that even if transcripts from both alleles are detected at the population level, there may be substantial variation in the degree of allelic imbalance in single cells. For example, while for most genes individual cells express RNA from both alleles, for other genes the population can be a mixture of cells expressing RNA from only one or the other allele. This latter expression pattern has been termed random monoallelic expression, and certainly, genes with such an expression profile exist: most X-linked genes are expressed only from one X chromosome due to random X-chromosome inactivation (X. Deng et al. 2014; Galupa and Heard 2015; Payer 2016), and a similar pattern has also been shown for some autosomal genes, such as olfactory receptors or antigen receptors (Monahan and Lomvardas 2015; Brady, Steinel, and Bassing 2010; Eckersley-Maslin and Spector 2014). Understanding random monoallelic expression is of particular interest given that quantitative cell-to-cell differences or spatial heterogeneity in allele-specific gene expression have the potential to modify phenotypic outcome if the two alleles harbor different functional variants, as has been described for both X-linked (eg. (Yoshioka, Yorifuji, and Mituyoshi 1998; Plenge et al. 2002; Renault et al. 2007; Simmonds et al. 2014; Echevarria et al. 2016)) and autosomal traits (Pereira et al. 2003; Raslova et al. 2004).

Beyond these prototypic cases, it has been proposed that many more autosomal genes may be subject to random monoallelic expression (Gimelbrant et al. 2007; Zwemer et al. 2012; Li et al. 2012; Gendrel et al. 2014; Eckersley-Maslin et al. 2014; Q. Deng et al. 2014; Reinius et al. 2016), but some key properties of this extended class sets them apart from the more established examples. Similarly to the initial group, these genes were classified as displaying random monoallelic expression because cells with only transcripts from one or the other gene copy were observed, with both types of cells present in the same experiment. In addition, some of these genes maintain their monoallelic expression status over multiple passages in clonal cell lines (Gendrel et al. 2014; Reinius et al. 2016), so a specific, heritable mechanism could limit transcription to only one allele, as is the case for X chromosome inactivation. However, thus far such a mechanism remains elusive (Gendrel et al. 2014; Eckersley-Maslin et al. 2014). Moreover, unlike previously established cases, many genes in this extended set are not expressed exclusively monoallelically, and typically a subset of cells or clones with transcripts from both alleles can also be detected (Gimelbrant et al. 2007; Zwemer et al. 2012; Gendrel et al. 2014; Li et al. 2012; Eckersley-Maslin et al. 2014). This suggests that if a specific mechanism does exist to regulate monoallelic expression, it is limited to only a subset of cells.

To resolve this question of mechanism, Reinius et al proposed an elegant scenario (so-called dynamic random monoallelic expression), whereby the many genes with random monoallelic expression may arise not by differential cell- and allele-specific regulation, but instead monoallelic expression may arise by chance (Reinius and Sandberg 2015; Reinius et al. 2016). In this scenario infrequent transcriptional bursting would lead to cells that contain only RNA from one of the two gene copies. This observed monoallelic expression of mRNA would be temporary and the allelic state of a cell could change over time. The authors confirmed this model in clonal cell lines (Reinius et al. 2016), but whether the same is true in tissues is still an open question. Different groups have deployed single-cell transcriptomics to determine the degree of cell-to-cell allelic imbalance (Q. Deng et al. 2014; Kim et al. 2015; Reinius et al. 2016; Jiang, Zhang, and Li 2017), but technical limitations inherent to low abundance RNA quantification, as well as parameter choice can impact the interpretation of allele-specific sequencing data. Thus, it has been hypothesised that the level of monoallelic expression, especially at single-cell level, could be an overestimate (DeVeale, van der Kooy, and Babak 2012; Kim et al. 2015). This absence of precise quantitative data has made it difficult to definitively answer if random monoallelic expression observed *in vivo* requires a dedicated mechanism or if it could arise as a consequence of transcriptional bursting.

In this study, we adapted a previously described single-molecule RNA fluorescent *in situ* hybridization technique that is sensitive to single-nucleotide differences between RNAs for the analysis of transcripts in snap-frozen, cryosectioned tissues from different mouse strains and their hybrids ((Levesque et al. 2013; Shaffer et al. 2015; Ginart et al. 2016)). This allowed us to determine the allelic origin of individual mRNAs in single cells, while preserving both their spatial context and their *in vivo* expression levels. We used this method to measure allele-specific expression of multiple autosomal genes and of *Xist*, a gene for which it is well-documented that individual cells randomly and exclusively transcribe either the maternal or the paternal copy. Quantitative analysis of the data enabled us to answer whether the autosomal genes we investigated were expressed from one or both gene copies in single cells in tissue. While we observed monoallelic expression in some cells, mathematical modeling showed that this pattern was compatible with random transcriptional events (including transcriptional bursting) from the two alleles producing low levels of RNA, rather than an explicit mechanism governing random monoallelic expression.

## Results

### SNV-specific detection of RNAs in mouse tissue using single molecule RNA FISH

Our goal in tissues was to quantitatively measure the amount of cell-to-cell variability in transcript abundance from either the maternal or paternal allele of a gene to determine the degree of imbalance between transcripts arising from the two alleles. To make these measurements, we modified a protocol previously developed by our group (Levesque et al. 2013) that enables the detection of single nucleotide variants (SNVs) on individual RNA molecules *in situ* in cultured cells. This method works by first detecting the mRNA of interest (regardless of the allele of origin) using conventional single molecule RNA FISH probes labelled in one color (guide probes), and then colocalizing this signal with probes that discriminate specific single-nucleotide differences based on a “toehold probe” strategy and which are labelled in colors unique to the two different alleles (Figure 1A). In this way, mRNAs are essentially “tagged” as being either from one or the other parental chromosome. In cultured cells, this approach can successfully distinguish RNA variants that contain just one single nucleotide variant (SNV) and thus can only be targeted with a single variant-specific probe (Shaffer et al. 2015; Levesque et al. 2013), but the decreased signal-to-noise ratio makes reliable detection of single probes more difficult in tissue (Richardson and Lichtman 2015). We therefore opted to work with C57BL/6J (BL6) and JF1/Ms (JF1) mice, which belong to two different *Mus musculus* subspecies (*domesticus* and *molossinus*, respectively) (Koide et al. 1998; Takada et al. 2013). Due to their distant relationship, JF1 mice harbor a large number (>50,000) of SNVs in genic regions compared to BL6 (Takada et al. 2013), allowing us to target most genes with multiple SNV-specific probes.

**Figure 1.**
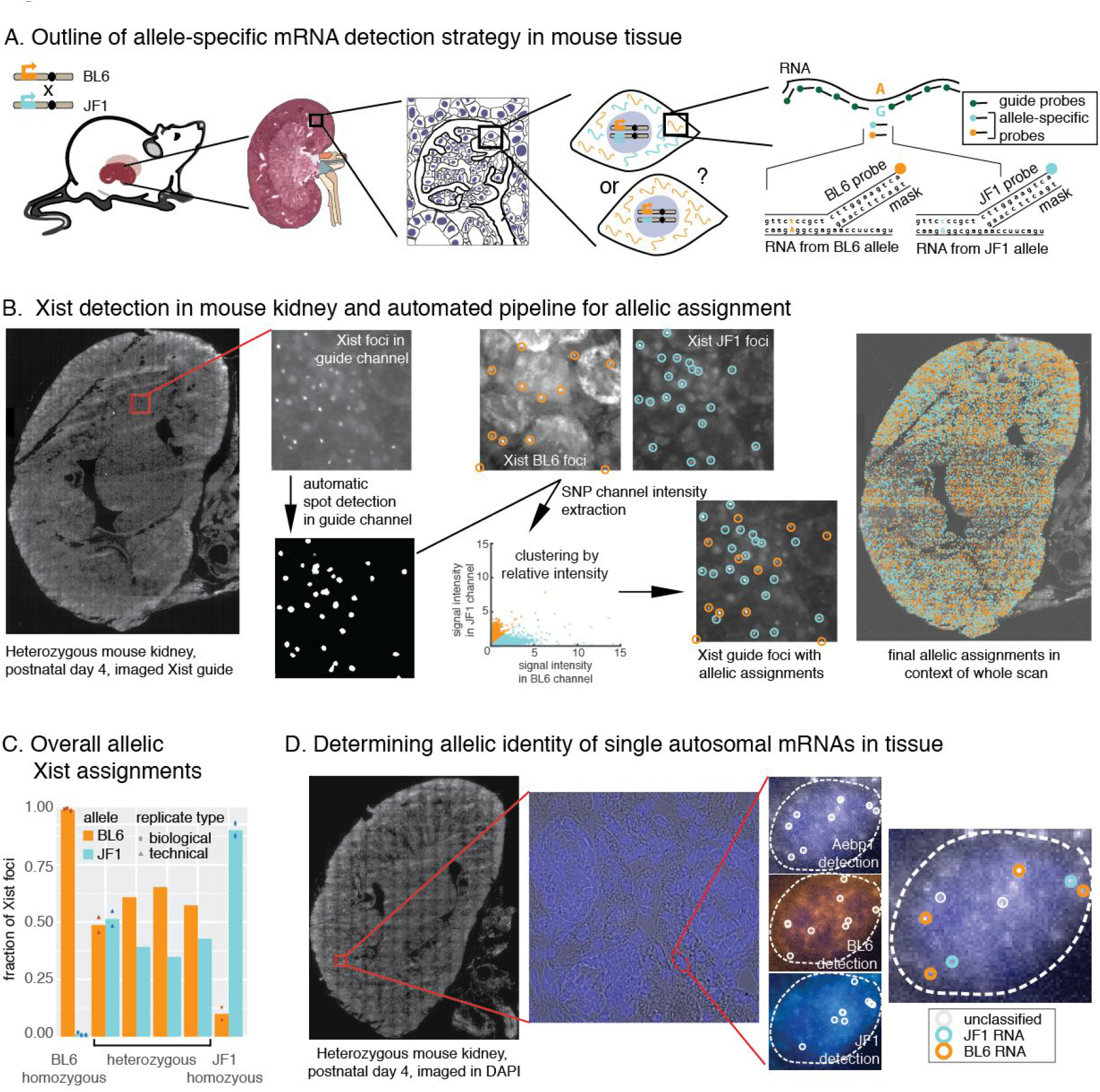
SNV FISH enables allele-specific detection of RNA in tissue. **A.** Collecting tissue from F1 heterozygous mice from crosses between BL6 and JF1 mice enabled allele-specific RNA detection in single-cells by utilizing a “toehold probe” strategy that targets polymorphisms between the two strains. While only a single polymorphism-probe is shown here for illustration, all genes were targeted with multiple SNV probes to increase the fluorescent signal. **B.** The pipeline for allele-specific detection of Xist foci in kidney tissue sections: First, whole-section tissue scans were used to collect fluorescence data in the guide and SNV channels (the scan shown is for the Xist guide probes labelled with Cal fluor 610). The guide probe signal was then used to automatically detect Xist foci, and the fluorescence intensity for the two allele-specific probes was extracted at the positions of those foci (BL6 was labelled with Cy3, JF1 labelled with Cy5). Clustering of the relative fluorescence intensities was applied to all foci, allowing automated allelic assignment for all spots across the entire tissue. **C.** Quantification of *Xist* allelic assignments in homozygous and heterozygous tissues. For each sample we depict the fraction of assigned BL6 and JF1 foci. For heterozygous samples biological replicates (ie tissue samples from different animals) are shown separately to reveal inter-individual variability in random X inactivation, while data from a technical replicate (two separate tissue sections from the same animal) is also plotted to show the effect of intra-individual variability and technical noise on allelic assignments. **D.** The pipeline for allele-specific detection of punctate mRNA spots in kidney tissue sections: whole tissue scans at 60x resolution in DAPI (staining nuclei) were used to obtain an overall impression of tissue morphology. Then, we collected z-stacks for randomly selected locations at 100x resolution in all channels of interest, which enabled detection of single cells and thresholding of Cal Fluor 610 detecting Aebp1 mRNA (top), BL6-detection probes labeled with Cy3 (middle), and JF1-detection probes labeled with Cy5 (bottom). Testing colocalization of these probes revealed allelic assignments for guide probes (far right).

To demonstrate our ability to detect expression from the BL6 vs. JF1 allele using our technique in tissue, we first examined the expression of *Xist*, a prototypical example of random monoallelic expression (Sado and Brockdorff 2013; Galupa and Heard 2015). In female cells *Xist* is transcribed exclusively from the (randomly chosen) inactivated X chromosome, is expressed at high levels, and contains a large number of SNVs between the two substrains, making it an ideal test case for our method. We collected kidney tissues from mice at day 4 of postnatal development, snap-froze them in liquid nitrogen or on aluminium blocks in dry ice and cryosectioned them at 5 μm thickness. We then applied our *in situ* hybridization protocol using allele-specific probes. As expected, we observed the appropriate gender-specific expression pattern: no fluorescent signal in male tissues (Figure S1), whereas in female tissues, our guide probes clearly labelled nuclear *Xist* foci, which colocalized with signal from the strain-specific probes. For SNV-targeting probes, controls in homozygous tissues confirmed that only the appropriate probes bound, with little binding from the probes targeting the other strain (Figure S1). Whereas in heterozygous samples the two strain-specific probes each labelled a subset of the nuclei in the anticipated mutually exclusive pattern (Figure 1B). To further analyse the data, we developed a pipeline to computationally identify Xist RNA foci, and automatically classify them as BL6 or JF1 within an entire scanned kidney section (Methods, Figure 1B). This algorithm identified tens of thousands of Xist foci per section (mean 22k, min 8.2k, max 30k), and—in agreement with our manual inspection of the data—predominantly identified Xist foci of the correct identity in the homozygous samples (Table S1), while the overall population ratio of BL6:JF1 foci ranged from 45:55 to 65:35 in heterozygous samples (Figure 1C, Figure S2A).

Having verified that we could correctly measure the allelic origin of clusters of Xist RNA accumulated on the X chromosome in mouse tissues, we next ascertained whether the method would also work for monodisperse spots corresponding to single RNA molecules, which is how most mRNAs appear in the cell. We considered this challenging, because single, punctate RNA spots would be both smaller and considerably dimmer compared to Xist RNA, which accumulates multiple copies on the inactivated X chromosome. Thus, to test our protocol for use on single RNA molecules, we designed probes for 8 autosomal genes that contained at least one polymorphism in BL6 versus either JF1 or the C7 strain (which carries both copies of chromosome 7 from the *M. musculus castaneus* strain in a BL6 background). These genes were selected to represent genes with (*Aebp1, Churc1, Lyplal1*) and without (*Aqp11, Mpp5, Podxl, Prcp, Stard5*) putative random monoallelic expression (Gendrel et al. 2014; Eckersley-Maslin et al. 2014; Li et al. 2012; Zwemer et al. 2012) and with different expression levels, patterns and chromosomal locations. Some of the chosen genes were also linked to specific kidney disease phenotypes (*Aqp11 (Ishibashi, Hara, and Kondo 2009), Mpp5 (Straight et al. 2004; Weide et al. 2017), Prcp (Maier et al. 2017)*) (Table S4). As expected, these genes were typically expressed at much lower levels than *Xist* and punctate individual mRNA spots could not be as readily observed in low magnification scans. We therefore combined whole tissue scans in a single plane at low magnification with random sampling of the tissue at higher magnification, where we imaged the entirety of the section. This approach allowed us to identify individual mRNA spots within the context of the whole tissue section in the 60x scan, while the additional data collected from the 100x z-stacks facilitated our ability to precisely determine colocalization between the guide probes and the strain-specific probes (Figure 1D). Colocalization between the strain-specific signal and the mRNA probes allowed us to determine from which allele a given mRNA originated. Accordingly, this could be used as a key readout for SNV-specific single molecule RNA FISH, and we characterized the quality of the experiments using colocalization rate (*i.e*. what percentage of the guide spots colocalized with allele-specific spots). When we assayed the overall colocalization rates for these autosomal genes in kidney sections, we found that 4 out of 8 genes had mean colocalization rates >50% (Figure S3, Table S4), which is comparable to colocalization rates previously observed in cultured cells (Levesque et al. 2013), showing that we were able to perform quantitative SNV-specific single molecule RNA FISH directly in tissue. We also observed an apparent trend between colocalization rate and the number of SNV probes, where genes with fewer SNV probes had lower colocalization rates than those with more SNV probes. We tested a series of parameters that could affect probe binding (base composition, GC content, probe secondary structure and folding energy), but found no parameter that differentiated between the probes with high and low colocalization rates (Figure S3). However, it should be noted that SNV probes were only tested as full sets (*i.e*. all SNV probes for a given gene were tested together), and we therefore do not know the binding behaviour of individual probes.

Collectively, these results showed that we could directly visualise and assign strain-specific identities to both focally localised RNA (Xist) and single molecule RNA spots in the context of whole tissue sections. This motivated us to ask if we could directly quantify cell-to-cell heterogeneity in the allelic origin of RNAs in tissue.

### Quantifying cell-to-cell heterogeneity in the chromosomal origin of RNA in tissue

To determine how the chromosomal origin of RNAs contributes to cell-to-cell heterogeneity in tissue, we focused on two different questions. For *Xist*, we investigated the spatial clustering of cells based on their X inactivation choice, *i.e*. to what extent cells expressing Xist from either the JF1 *vs* BL6 X chromosome intermix. For the autosomal genes, we quantified their allelic imbalance in single cells, *i.e*. whether individual cells expressed RNAs from the two chromosomes at different ratios.

In the case of Xist, we measured spatial clustering of allele-specific expression because each cell is randomly and fully committed to expressing RNA from only the BL6 or the JF1 chromosome. Thus, allelic imbalance in tissue is not due to quantitative expression differences between the two alleles in single cells. Instead, it can arise either as a consequence of overall skewing of X chromosome inactivation rates or due to uneven spatial distribution of cells with a given inactivated X chromosome. Such spatial partitioning can arise through the local expansion of cells in which X chromosome inactivation “choice” has already been fixed, resulting in extended patches of cells carrying the same inactive X chromosome (Gardner et al. 1985; Ponder et al. 1985; Mrozek et al. 1991; Wu et al. 2014). Because we had observed fairly balanced expression from the BL6 and JF1 chromosome in our initial analysis of heterozygous samples (see previous section, Figure 1C), we next determined whether cells expressing Xist from either the BL6 or JF1 allele segregated spatially. Although visual inspection of the sections revealed no extended regions where cells expressed Xist foci with the same strain-specific identity, computational modelling could potentially reveal a more precise view of spatial patterning. We therefore developed a metric that characterized the distribution of cells expressing BL6 Xist RNA (Methods and Figure S2) and then compared this to either completely randomized BL6 and JF1 assignments or randomizations where we introduced different sized clusters of cells. We found that in all tissues BL6 cells were less evenly distributed than the random assignments (Figure S2C), suggesting that cells cluster together more than expected in a completely random scenario. Our subsequent comparison with the clustered assignments further supported and refined this interpretation: it showed that our data was most similar to simulations with smaller cluster sizes. The closest matching seed size was different, depending on what scale we assessed spatial partitioning, but centered around a cluster size of 2–4 (mean cluster size: 3.2, standard deviation 0.7) (Table S2, Figure 2A and Figure S2D). Thus, cells expressing *Xist* from the same chromosome clustered together in small patches in tissue sections, resulting in a spatially fairly mixed population of cells and showed no evidence of extended patches with the same allele-specific expression.

**Figure 2.**
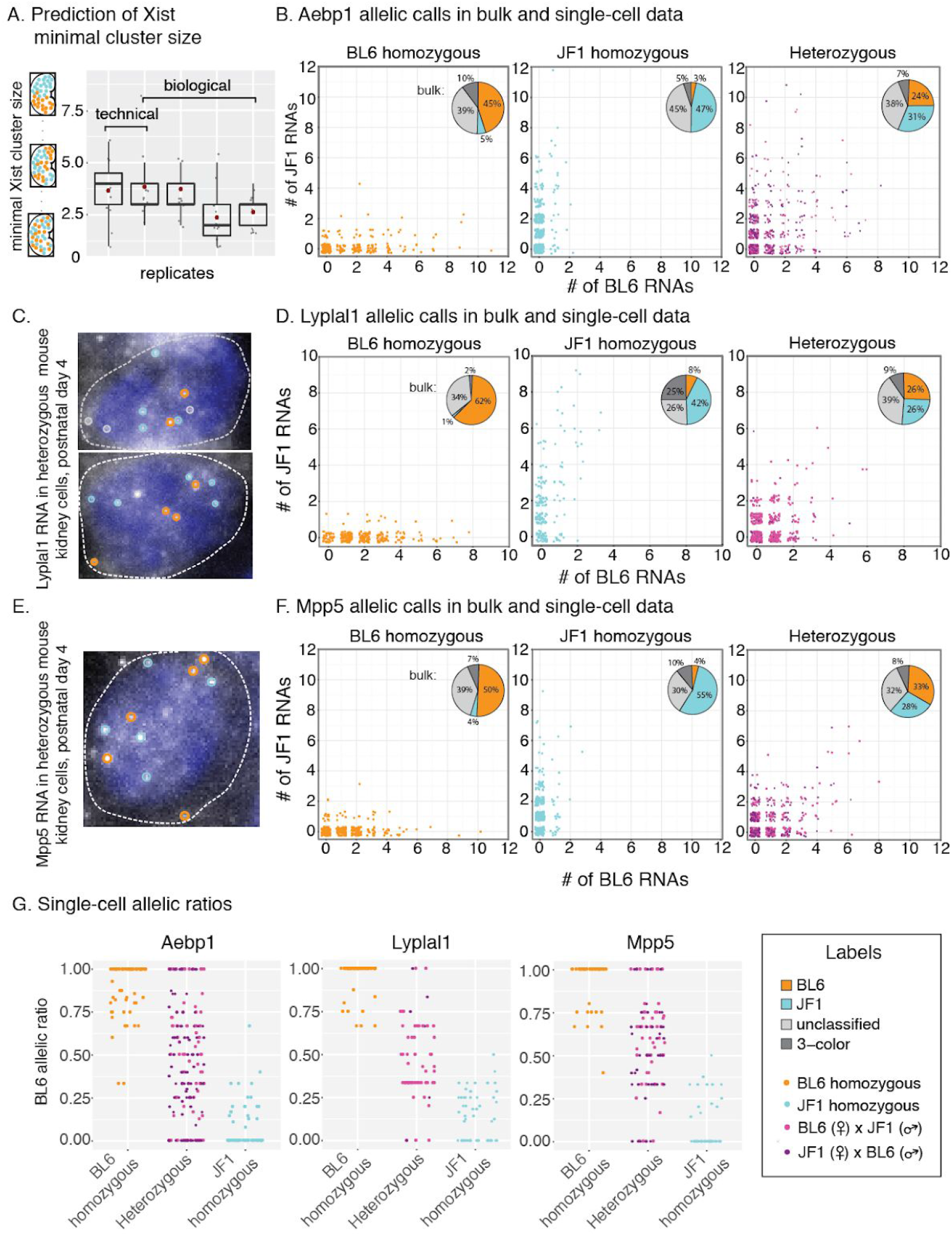
Quantification of allele-specific single-cell heterogeneity. **A.** Prediction of minimal size of Xist clusters with the same allelic identity. For each sample (x axis) we show the seed size with the closest matching variance for different subdivision sizes (grey dots, for original data see Figure S3). The overlaid box plots show the overall distribution of these data points, with the red dot indicating the mean seed size from all subdivision sizes. **B,D and F.** Measurements of allele-specific expression in single cells (scatter plots) and in bulk data (pie charts) for Aebp1 (B), Lyplal1 (D) and Mpp5 (F). For each gene, data are shown for BL6 homozygous (left), JF1 homozygous (middle) and heterozygous (right) samples. For the single-cell scatter plots cells only contained integer numbers of RNA, but we included jitter to better display the density of cells with a given allelic distribution. For all plots we combined data from different replicates with more than 40% colocalization rate (summaries for each experiment are shown in Supplementary Table 5), and the two colors in the scatter plots indicate data from the two reciprocal crosses (ie BL6 x JF1 and JF1 x BL6). **C and E.** Fluorescence micrographs and allelic assignments of Lyplal1 (C) and Mpp5 (E) guide spots in BL6 x JF1 heterozygous kidney cells. For both genes guide probes were labelled with Cal fluor610. **F.** BL6 allelic ratios in single cells with >2 RNA with allelic assignments for homozygous and heterozygous samples for Aebp1 (left), Lyplal1 (middle) and Mpp5 (right).

For the autosomal genes we wanted to know how much the allelic imbalance differed between cells. Visual inspection of our data indicated a range of allelic ratios, including BL6 monoallelic, JF1 monoallelic, as well as biallelic mRNA expression. To quantify this, we focused on three autosomal genes (*Aebp1, Lyplal1, Mpp5*) where >50% of guide spots colocalized with signal from SNV probes (we excluded *Podxl*, because the high expression levels of Podxl mRNA in podocytes precluded separating individual RNA spots (Figure S4)). First, we considered that the observed chromosomal origin (BL6 vs. JF1) could either be due to true biological variability or to technical error (as seen when we detected RNA from the “incorrect” strain in homozygous tissues). To distinguish these, we determined the BL6 and JF1 signal for these genes in kidney tissue from both BL6 and JF1 homozygous mice, as well as in tissue from reciprocal heterozygous crosses. For all three genes, we counted only a few mRNAs in the majority of cells (mean number of RNA spot counts per cell: 3.5 for *Aebp1*, 3.2 for *Lyplal1* and 2.6 for *Mpp5*), and we predominantly detected the correct allele in the homozygous kidney samples, both in bulk and at the single cell level (Figures 2B, D and F). In heterozygous samples we observed a more balanced presence of both BL6 and JF1 mRNAs, with the reciprocal crosses showing similar results (heterozygous data in Figures 2B, D, F and G, and Table S5). These results indicated that our technical error (false positive rate) was less than the biological variability and that we could use our method to measure quantitative single-cell differences. Moreover, when we compared the BL6 allelic ratios in homozygous and heterozygous cells with >2 mRNA, we found some heterozygous cells with allelic ratios similar to those of the homozygous samples, but also a subset of cells with an allelic ratio that was intermediate to that of homozygous cells (Figure 2G). These results were particularly intriguing for *Aebp1* and *Lyplal1*, because these two genes had been previously identified as genes with putative random monoallelic expression in other tissues (Zwemer et al. 2012; Li et al. 2012; Gendrel et al. 2014; Eckersley-Maslin et al. 2014). Given that we observed cells with either BL6 or JF1 monoallelic mRNA expression in heterozygous tissue, akin to the random monoallelic expression pattern, we wanted to know if we could use our quantitative single-cell allelic imbalance data to explain how this expression patterns could arise.

### Observed heterogeneity of strain-specific RNA is compatible with bursty biallelic expression

Our results showing that individual cells could have mRNA from either one or both alleles motivated us to assess in more detail whether existing models of transcription were sufficient to explain the observed cell-to-cell variability in allelic imbalance. For this, we initially considered two extreme cases: an “all-or-none” scenario in which every cell has transcripts exclusively from one or the other gene copy, and a “coin flip” scenario in which the allelic origin of every individual transcript is essentially indistinguishable from random coin flipping (Figure 3A). The former scenario could suggest the existence of regulatory mechanisms that limit transcription to only one gene copy per cell (an extreme form of random monoallelic expression, as is the case with *Xist*), whereas the latter corresponds to a null model with no distinct allele-specific transcriptional regulation. We used computational modeling to simulate these two scenarios and to discriminate between these two scenarios.

**Figure 3.**
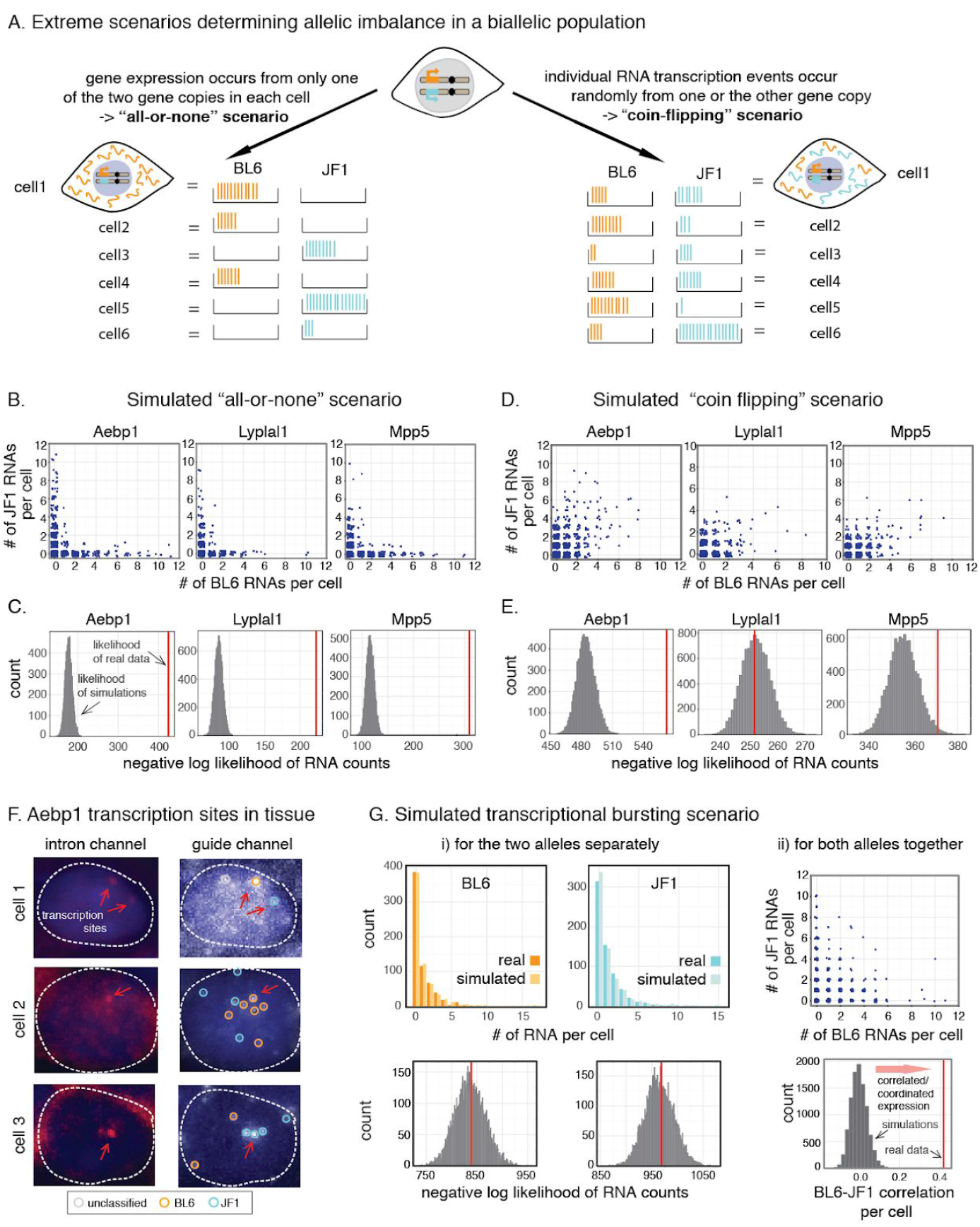
Determining the mechanisms underlying allele-specific single-cell heterogeneity. **A.** Outline of two extreme scenarios that could lead to variable single-cells allelic imbalance. **B, C.** Simulated single-cell RNA counts (B) and negative log likelihood distribution of simulated and observed data (C) in the case of the “all-or-none” model. **D, E.** Simulated single-cell RNA counts (D) and negative log likelihood distribution of simulated and observed data (E) in the case of the coin-flipping model. **F.** Single-molecule RNA FISH of *Aebp1* introns (left) and guide spots (right) in mouse kidney, including a cell with two transcription sites (top), a transcription site on the BL6 allele (middle) and a transcription site on the JF1 allele (bottom). The position of the transcription sites are highlighted by red arrows and the allelic assignment of the guide spots are indicated by colored circles. Introns were labelled with Atto488 (top) or Alexa700 (middle, bottom). **G.** Outcome of simulating the “transcriptional bursting” scenario for each allele separately (i) and for the alleles together (ii). Top row represents simulated single-cell RNA counts, bottom row shows negative log likelihood distribution of simulated and observed data for the BL6 and JF1 allele individually (i) and the correlation between randomly paired simulated RNA counts (ii). All log likelihood distributions (in C, E and G) and the correlation distribution (in G) were obtained from 10,000 simulations, and one of these simulations was picked randomly for each gene to display RNA distributions in B, D and G.

We first checked if our data were similar to those expected in the “all-or-none” scenario, in which the transcripts in each individual cell were either solely from one or the other gene copy. Looking in heterozygous cells, it seemed qualitatively apparent that (as noted) many cells have mRNAs from both gene copies, which was seemingly incompatible with the all-or-none scenario (heterozygous data Figure 2B, D, F and G). However, it was still formally possible that cells in reality only had transcripts exclusively from either one or the other gene copy and that the apparent transcripts from the other copy were technical artifacts due to false detection events. We used homozygous tissue to measure the rate at which these false detection events occur, and thereby estimated the expected false detection rate in heterozygous cells. We then computationally simulated a hypothetical “all-or-nothing” heterozygous cell populations taking into account these false detection rates, and found that our actual data was still qualitatively inconsistent with the all-or-none hypothesis (compare Figure 3B with heterozygous data in Figure 2B, D and F). Moreover, we calculated the probability of the observed strain-specific RNA counts in the real *versus* the simulated cell populations, which revealed that the likelihood of the measured data was well outside that of the simulated distribution of likelihoods (Figure 3C). Collectively, these results indicate that in the kidney, none of the three genes we interrogated displayed “all-or-none” expression.

At the other extreme, it is possible that the two copies of the gene transcribe RNA independently, with each random transcription event producing a single RNA, thus leading to the “coin-flipping” scenario in which most cells would have RNAs from both alleles in them, but with some statistical noise about this population average. We modeled the outcome of such a scenario and found that the real vs. simulated single-cell RNA distributions looked very similar for all three genes (compare Figure 3D with heterozygous data in Figure 2B, D and F). By treating the strain-specific RNA counts per cell as a series of independent coin-flips, we then calculated the probability of the observed distributions. We found that for both *Lyplal1* and *Mpp5* the likelihoods of the real population measurements fell within the distribution of likelihoods from our simulated model (Figure 3E).

These results indicated that the allelic imbalance observed for *Lyplal1* and *Mpp5* was compatible with a simple coin-flipping null model of transcription from the two alleles, but *Aebp1* showed higher levels of imbalance per cell than could be explained by this model. Yet this gene did not exhibit the all-or-none behavior either. We therefore considered a third, intermediate scenario motivated by the phenomenon of transcriptional bursting (Raj and van Oudenaarden 2008; Vera et al. 2016; Nicolas, Phillips, and Naef 2017). Transcriptional bursts refer to the fact that most mammalian genes are transcribed in short pulses during which multiple transcripts are synthesized, interspersed between periods during which the gene remains inactive. When there are two copies of a gene, each bursting independently (Levesque and Raj 2013), the expected result would be that some cells may have more transcripts from one of the copies than expected by the coin flipping model above; bursting would be akin to getting several heads or tails in a row every time one flipped a coin.

To test whether transcriptional bursting could explain the observed data, we first wanted to confirm that *Aebp1* was indeed transcribed in a burst-like fashion. To verify this, we measured *Aebp1* transcriptional activity directly in kidney cells by using intronic probes (Figure 3F), which, owing to the extremely short half-life of introns, detect almost exclusively nascent transcripts at the site of transcription (Levesque and Raj 2013). This showed that 19% of cells with Aebp1 mRNA were also actively transcribing *Aebp1*, and that these transcription sites contained more than 1 RNA based on their fluorescence intensity relative to cytoplasmic RNA spots (average 1.6x higher fluorescence intensity in 3 independent experiments, Figure S5). This data also showed that the majority of actively transcribing cells had only one *Aebp1* transcription site, although a small subset (6 out of 55 (11%) cells with *Aebp1* transcription sites) showed simultaneous expression from both alleles. This corroborated that cells indeed produced *Aebp1* in transcriptional bursts and that it was possible for individual cells to transcribe RNA from both alleles simultaneously.

Next, we turned to simulations to assess the RNA distributions that we would expect in a scenario where the two alleles transcribed RNA independently from each other and in bursts. Initially, we estimated the expected burst size (average number of RNAs that were transcribed together in a single burst) and burst frequency for the two alleles based on the observed RNA counts for each allele independently and used these parameters as inputs for our model (see methods for details). BL6 and JF1 RNA counts simulated this way closely matched our measurements, which was also reflected by the likelihood of the real data falling within that of the simulated data for the two alleles separately (Figure 3G). We then wanted to see whether the degree to which there was allelic imbalance in single cells could be explained by the two alleles bursting independently; *i.e*., whether for per-cell RNA counts from the two alleles was more or less correlated than one would expect by chance. To simulate the null hypothesis of no interaction between alleles we randomly paired up the modeled BL6 and JF1 counts, mimicking cells that contain RNA from both alleles. When we compared this simulation to the real data we consistently observed that the modeled per cell BL6 and JF1 counts were were less correlated that the real pairwise measurements (Figure 3G and Figure S6). Thus, while in our measurements cells with high BL6 expression typically also expressed JF1 at higher levels (compare heterozygous data in Figure 2B and 3G top panel), in the modeled data BL6 and JF1 counts showed little correlation. This was also true when we incorporated false detection events in our model to account for possible incorrect allelic assignment, as we had done in the “all-or-none” model (Figure S6). Together, our transcription site measurements and simulations showed that the observed allele-specific single-cell RNA counts for *Aebp1* were compatible with transcriptional bursting of the two gene copies individually and that expression from the two alleles was correlated.

Thus, for *Aebp1, Lyplal1* and *Mpp5* the observed monoallelic expression in some single cells can likely be explained by low levels of transcription occurring randomly from the two gene copies without having to invoke a special mechanism that limits expression to one of the alleles.

## Discussion

There has been great interest in recent years to precisely measure expression from the two alleles of a gene in diploid cells, ideally directly in tissue and at the single-cell level. The RNA fluorescence *in situ* hybridization method described here is a step in this direction: by visualizing endogenous SNVs it enables the assignment of single RNA molecules to their allele of origin in single cells and in the context of whole tissue sections. We now provide quantitative information about cell-to-cell heterogeneity with single transcript resolution, which is an extension of our previous work, where we used this method for a more qualitative assessments (*i.e*. presence-absence) of parental origin of mRNA in tissue (Ginart et al. 2016). When we applied our method to autosomal genes we observed that individual cells in heterozygous tissue spanned the entire range from all RNAs originating from the BL6 chromosome through various more mixed populations to all RNAs originating from the JF1 chromosome. These observed allelic imbalances were not due to the parental origin of the gene copies, because reciprocal crosses (BL6×JF1 vs JF1×BL6) showed similar results. We therefore asked what model could explain the observed single-cell allelic imbalance pattern and combined our data with computational simulations to address this question. We found that the observed allelic distributions could be recapitulated by a model where transcription occurred randomly from the two alleles, perhaps with moderate transcriptional bursting (*e.g*. in the case *of Aebp1*). Thus, we did not have to invoke a special mechanism that restricted expression to only one allele to explain the presence of cells with either BL6 or JF1 monoallelic expression status.

Our results suggest that cells with RNA expression from only one of the alleles occur due to the low levels of expression and thus the limited number of random sampling events from the two gene copies. Because transcription is a dynamic process this state is likely transient so that while a cell may have mRNA from only one allele at a particular time, it may gain mRNA from the other allele a short time later if another transcriptional event occurs. For *Mpp5*, this conclusion is in line with the fact that the reported haploinsufficient phenotype is thought to be caused by overall reduced dosage (Straight et al. 2004; Weide et al. 2017) rather than by a special subpopulation of cells with monoallelic expression. More generally, this scenario links the observation of transcriptional bursting with that of random monoallelic expression, as put forward by Sandberg et al (Reinius and Sandberg 2015; Reinius et al. 2016), and explains why we (and others previously (Gendrel et al. 2014; Eckersley-Maslin et al. 2014)) observed the co-occurrence of cells with one and two transcription sites in the same population. Similarly, it could explain why many genes with monoallelic expression in one clonal cell line are expressed biallelically in others (Gimelbrant et al. 2007; Zwemer et al. 2012; Li et al. 2012; Gendrel et al. 2014; Eckersley-Maslin et al. 2014) and why the expression of a given gene is often lower in cell lines with monoallelic expression than in those with biallelic expression (Li et al. 2012; Zwemer et al. 2012; Gendrel et al. 2014). In the case of *Aebp1* and *Lyplal1*, which had previously been identified to display random monoallelic expression, our data suggest that no additional mechanism is needed to explain the presence of cells with monoallelic expression, but we cannot exclude the possibility of allele-specific regulatory mechanisms or maintenance of allelic status, especially in different mouse strains and different cell types that were used in the original studies.

In addition to the data on *Aebp1, Mpp5* and *Lyplal1*, we also demonstrated our ability to distinguish *Xist* expression from the BL6 vs JF1 chromosome and to assess the spatial relationship of cells expressing different parental alleles. We detected a spatially fairly mixed population in the kidney, where cells expressing *Xist* from the same chromosome clustered together in small patches in transverse sections. This is contrary to other tissues, such as intestinal crypts or the skin (Ponder et al. 1985; Thomas, Wiliams, and Wiliams 1988; Gardner et al. 1985; Wu et al. 2014), and suggests either a larger number of kidney precursor cells or extensive cell migration during development. Because our method only provides a snapshot in time we cannot easily distinguish between these scenarios (lineage vs. migration), especially given that the kidney is a complex organ, composed of cells originating from different embryonic lineages that undergo extensive migration during development even after birth (Takasato and Little 2015; Little and McMahon 2012). Regardless of the developmental mechanism, however, our data indicate that there is likely no major spatial segregation due to X chromosome inactivation in this tissue, and in the case of mutations, any phenotypic effects would be fairly evenly distributed.

Through the examples detailed above we have shown how our method can be used to directly quantify cell-to-cell differences that arise due to differential expression from the two alleles in diploid cells. Our approach overcomes multiple limitations imposed by previous methods: First, because this method enables sensitive SNV-specific detection of even single mRNA molecules it provides more information than RNA FISH measurements that rely solely on quantifying the number of transcription sites in individual cells (Raslova et al. 2004; Gimelbrant et al. 2007; Eckersley-Maslin et al. 2014; Gendrel et al. 2014; Huang et al. 2017). Second, although our approach tests only single genes in a given experiment and thus has much lower throughput than single-cell sequencing-based methods, it relies on direct detection of transcripts and is therefore not subject to subsampling and dropout, which complicate the interpretation of sequencing-based cell-to-cell variability results (Brennecke et al. 2013; Reinius and Sandberg 2015; Dueck et al. 2016; Jiang, Zhang, and Li 2017; Torre et al. 2018). Finally, by making use of pre-existing endogenous SNVs it eliminates the need for genetic manipulation, for example to label the gene of interest with fluorescent tags, as has been done to measure X chromosome inactivation choice (Wu et al. 2014; Kobayashi et al. 2016) or to monitor the transcription of autosomal genes from the two chromosomes (Aseem et al. 2013; Fritzsch et al. 2018). Moreover, because the breeding history for classical inbred mice has lead to extended regions of shared ancestry (and shared SNVs) between different strains (Frazer et al. 2007; Yang et al. 2011), a probeset developed for one strain can often easily be adapted for another strain. For example, while we measured *Xist* expression from the BL6 and JF1 allele, the probes were designed so that they should distinguish equally well between the 129-strains and CAST/EiJ, which are also commonly used to study strain-specific expression.

In addition, while our quantitative analysis focused solely on genes with a relatively high number of SNVs and high mean colocalization rates (>50%), it should be noted that we did not systematically explore the relationship between SNVs and colocalization rates, and also that lower colocalization can be sufficient to address specific questions, as was demonstrated in a recent single cell *in situ* analysis of A-to-I RNA editing (Mellis et al. 2017). It is therefore likely that depending on the scientific question, less stringent cutoffs can be applied to colocalization rates and/or the number of SNVs required.

In conclusion, we demonstrated how quantitative measurement of allele-specific expression in tissue could be used to directly determine the level of allelic imbalance in single cells. By combining these measurements with modeling, we showed that random monoallelic expression could arise *in vivo* by chance alone. Beyond this application, our methods could have a number of additional uses. Similar analyses could be performed in other tissues and, for example, could enable the evaluation of genetic variants directly in the tissue believed to be affected if there are genic SNVs in linkage with those variants or to study mutations thought to lead to haploinsufficiency. Furthermore, with single cell resolution, our method allows for the interrogation of particular cellular subtypes within a tissue. In concert with recent genome-wide association studies in single cells (Wills et al. 2013; van der Wijst et al. 2018), this technique provides a useful tool for quantitative assessment of allele-specific genetic effects.

## Methods

### Mice, tissue harvest, sectioning and fixation

C57BL/6J and JF1/Ms founder mice were obtained from Jackson Laboratories. All mouse work was conducted in accordance with the University of Pennsylvania Institutional Animal Care and Use Committee. For tissue collection we used either homozygous C57BL/6J or JF1/Ms pups, or F1 heterozygotes from both C57BL/6J x JF1/Ms or JF1/Ms x C57BL/6J crosses. We dissected pups at postnatal day 4 using standard techniques, and mounted tissues in Tissue-Plus O.C.T. compound (Fisher Healthcare), flash-froze them in liquid nitrogen or on an aluminum block in dry ice, and then stored tissues at −80°C. We determined sex of the animals by visual inspection and verified this by SRY-specific PCR on DNA extracted from a tail sample, collected during dissection. Tissues were cryosectioned at 5 μm using a Leica CM1950 cryostat. We adhered tissue samples to positively charged Colorfrost plus slides (Fisher Scientific), washed slides in PBS, fixed them in 4% formaldehyde for 10 min at room temperature, then washed again in PBS two times. Fixed slides were stored in 70% ethanol at 4°C.

### Probe design, synthesis and labelling

To identify exonic SNVs between the C57BL/6J and JF1/Ms strains we used the NIG Mouse Genome Database (http://molossinus.lab.nig.ac.jp/msmdb/index.jsp) (Takada et al. 2013). For *Aebp1* and *Lyplal1* we confirmed these SNVs through PCR amplification and sequencing of exonic sequences of genomic JF1/Ms DNA. All guide probes and the *Aebp1* intron probe set were designed using the Stellaris probe designer (Biosearch Technologies), SNV-specific probes were designed as specified in Levesque *et al*. (Levesque et al. 2013) and mask oligonucleotides were selected to leave a 7–11 bp overhang (toehold) sequence (all probe sequences available in Table S6). Guide probes were purchased labeled with Cal fluor 610 (Biosearch Technologies), while SNV-specific probes and intron probes were ordered with an amine group on the 3' end. For these latter probes we pooled the oligonucleotides for each probe set and coupled them to either NHS-Cy3 or NHS-Cy5 (GE Healthcare) for the allele-specific probesets, or NHS-Atto488 or NHS-Atto700 (Atto-Tec) for the intronic probes. We purified dye-coupled probes by high-performance liquid chromatography. Mask oligonucleotides were used unlabelled.

### DNA extraction, gene sequencing and sex-specific aenotvpina

DNA was extracted from tail biopsies using a quick-lyse protocol: 100μl of Solution A (25mM NaOH and 0.2mM EDTA) were added to the tissue and kept at 95°C for 30 min, before adding an equal volume of Solution B (40mM Tris, pH=8). Samples were then spun at 6000 rpm for 10 min and 100μl of the top layer was transferred to a fresh tube. 1μl of this solution was used as template for PCR. To verify the presence of reported SNVs in *Aebp1* and *Lyplal1*, we designed primers for the exonic segments of these genes (primer sequences available in Table S7), and PCR-amplified genomic DNA using AmpliTaq Gold (ThermoFisher) with buffer II and 0.25mM MgCl_2_ according to the manufacturer's instructions. PCR amplicons were purified with ExoSAP-IT (ThermoFisher) and submitted for sequencing to the University of Pennsylvania DNA sequencing facility. For sex-specific genotyping of pups we used *Sry*-specific primers (Table S7), since this gene is located on the Y chromosome and thus amplicons can only be detected in male tissues. PCR was performed as for sequencing, and the presence-absence of a product was revealed on a gel.

### Allele-specific RNA fluorescence in situ hybridization of tissues

For each gene of interest we first prepared a probe mix, containing a guide probe set (labelled with Cal fluor 610), the two allele-specific probe sets (labelled in Cy3 and Cy5, respectively) and a set of mask oligos (unlabelled, in 1.5x excess of the allele-specific probes) in hybridization buffer (10% dextran sulfate, 2× SSC, 10% formamide). For detection of nascent Aebp1 RNA we also included intronic probes labelled either with Atto488 or Atto700, and to verify the integrity of RNA in male tissues stained for Xist we also included *Gapdh* probes (labelled with Atto488). To stain the samples, we first washed the slides with tissues sections in 2x SSC, then incubated them in 8% SDS for 2 minat room temperature, washed again in 2x SSC and finally added the hybridization buffer with probes. Slides were covered with coverslips and left to hybridize overnight in a humidified chamber (ibidi) at 37°C. The next morning we performed two 30 min washes in wash buffer (2× SSC, 10% formamide), the second one including DAPI to stain nuclei. To label cell membranes (to clearly identify single cells) the first wash was sometimes substituted with a 15 min incubation in wash buffer containing wheat germ agglutinin coupled with Alexa488 (LifeTech) and a 15 min regular wash. After the final wash, slides were rinsed twice with 2x SSC and once with antifade buffer (10 mM Tris (pH 8.0), 2× SSC, 1% w/v glucose). Finally, slides were mounted for imaging in antifade buffer with catalase and glucose oxidase (Raj et al. 2008) to prevent photobleaching.

### Imaging

We imaged all samples on a Nikon Ti-E inverted fluorescence microscope using either a 60x or a 100× Plan-Apo objective and a cooled CCD camera (Andor iKon 934). For whole-tissue scans we imaged at 60x and used Metamorph imaging software (Scan Slide application) to acquire a tiled grid of images. We used the Nikon Perfect Focus System to ensure that the images remained in focus over the imaging area. For 100× imaging, we acquired *z*-stacks (0.3 μm spacing between stacks) of stained cells in six different fluorescence channels using filter sets for DAPI, Atto 488, Cy3, Calfluor 610, Cy5, and Atto 700. The filter sets we used were 31000v2 (Chroma), 41028 (Chroma), SP102v1 (Chroma), 17 SP104v2 (Chroma), and SP105 (Chroma) for DAPI, Atto 488, Cy3, Cy5, and Atto 700, respectively. A custom filter set was used for CalFluor610 (Omega).

### *Xist* image analysis and modelling

For Xist image analysis we worked with whole tissue scans, where we had collected data for Cal fluor 610 (Xist guide probes), Cy3 and Cy5 (BL6 and JF1 probes, respectively) and DAPI (nuclei). To visualize scans, we used the “Grid/Collection stitching” feature available in Fiji (Schindelin et al. 2012) to assemble tiles. To identify Xist RNA and assign them an allelic identity we developed a custom pipeline in matlab. First, we reconstructed the scan taking into account the tile order provided in a supplementary file. Then, we used the data from the guide channel to detect Xist foci, regardless of allelic identity: we performed background subtraction, removed small objects and smoothened boundaries by border clearing and morphological opening, and then used LoG filtering to sharpen objects, binarized the observed signals and created connected components. Visual inspection of these connected components showed that they largely corresponded to Xist foci, but some areas with high background signal were also being detected as connected components. We therefore applied a number of filters (minimum fluorescence intensities for all RNA FISH channels, minimum cutoff for solidity, maximum area for connected components) to yield the final segmentation. Each obtained spot was then parametrized as the ratio of the signal intensity (background subtracted and normalized to the mean intensity of the scan) of the two SNV probe channels and we applied k-means clustering (2 means) to yield a critical angle above which we assigned spots JF1 identity, and below which we assigned BL6 identity. To verify the quality of these assignments, we designed a graphic user interface to manually annotate Xist foci and their allelic identity. We typically annotated 10 or more randomly selected tiles and the results of this quality control step are shown in Table S3. On average ~90% (mean 90.9%, standard deviation 5%) of Xist foci were correctly detected, while the remaining 10% of identified spots were areas of high background intensity that had been miscategorized as Xist foci. When Xist spots were correctly identified, typically more than 90% were assigned the correct allelic identity (mean 94.4%, standard deviation 5%). To assess spatial patterns of Xist allelic choice we then used the positional and identity information from our automatic assignments, and developed a metric for spatial heterogeneity. First, we tiled images into regular rectangles of equal size (i.e. 16 tiles all 1/16 of the full scan size). For each rectangle, we calculated the fraction of cells expressing Xist from the BL6 allele. Next, we obtained the variance of these BL6 cell fraction values across all rectangles of a given size. This protocol was repeated for different sizes of rectangles ranging from 16 to 256 rectangles spanning the entire tissue section. We also calculated a baseline for spatial heterogeneity of random allelic choice by repeating this analysis on 1000 random permutations of the data for each sample generated by Matlab’s randperm. We performed a similar analysis to determine the minimal cluster size of Xist foci with identical allelic identity, but instead of random permutations we generated simulations, where kidney sections were randomly seeded with clusters of a fixed size (ranging from 1 to 10) while keeping the allelic ratio the same as for the measured data. For each seed size we generated 500 simulations. To obtain a likely minimal cluster size for cells with identical X chromosome inactivation we selected the seed size whose variance deviated least from the variance observed for the real data. We repeated this process for each subdivision size and determined the mean across all subdivision sizes.

### Allele-specific colocalization analysis of single mRNA spots

For analysis of single molecule RNA spots we used a combination of 60x whole tissue scans in DAPI and Cal fluor 610 to determine the overall structure of the tissue and collecting z-stacks at 100x resolution of 5–10 individual positions within that tissue to identify individual mRNA molecules and characterize their allelic identity. To determine allelic identity we first segmented and thresholded images using a custom Matlab software suite (downloadable at https://bitbucket.org/arjunrajlaboratory/rajlabimagetools/wiki/Home, changeset: d278b7d0012282ecb318fde3bebbe3beaba62032). To quantify colocalization rates we first determined the ideal colocalization radius for each gene. To do so, we segmented extended areas of the tissue (typically containing 10–50 cells). To ascertain subpixel-resolution spot locations the software then fitted each spot to a two-dimensional Gaussian profile specifically on the z plane on which the spot occured. Next, colocalization between guide spots and allele-specific spots was determined in two stages. In the first stage, we searched for the nearest-neighbor allele-specific probe for each guide spots within a 2.5-pixel (360-nm) window and ascertain the median displacement vector field, which was subsequently used to correct for chromatic aberrations. After this correction, we tested a range of different radii (r= 0.1 to 2.5 pixel) for each gene to calculate colocalization rates for the real data, as well for pixel-shifted data, where we took our images and shifted the guide channel by adding 2*r pixels to the X and Y coordinates. This pixelshifted data was used to test random colocalization due to spurious allele-specific spots. For each gene we then visually inspected colocalization rates for real and pixel-shifted data at the different radii and determined a radius where both the colocalization rate for the real data and the difference between the real and the pixel-shifted data was maximal. The selected colocalization radii for each gene are included in Table S4. To obtain allele-specific data for single cells we then repeated the colocalization analysis, but segmentation of cells was done by drawing a boundary around nonoverlapping individual cells using brightfield or wheat germ agglutinin signal, and colocalization was determined using only the previously determined ideal colocalization rate.

### Analysis of transcription sites

Transcription site analysis was performed using a custom Matlab software suite (downloadable at https://bitbucket.org/arjunrajlaboratory/rajlabimagetools/wiki/Home). For this, we segmented cells, thresholded RNA FISH signal and identified transcription sites for *Aebp1* by co-localization of spots in the intron and exon channel. Relative fluorescence intensities of transcription sites vs cytoplasmic RNAs were determined based on the fluorescence intensity of the guide probes using custom scripts written with R packages dplyr and ggplot2.

### Analysis of SNV orobe properties

To determine whether any biophysical properties could differentiate between allele-specific probes that had high vs low colocalization rates, we compiled a table containing the the following parameters (Table S8): probe name, probe sequence, colocalization rate (the colocalization rate determined for an entire probeset was applied to each individual probe), number of predicted secondary structures and folding energies. The latter two parameters were extracted by running sequences on the mfold web server (Zuker 2003) for DNA probes, with Na concentration set to 0.3M. Frequency of individual nucleotides, dAT, dGC, purines and pyrimidines was determined through analysis of the probe sequences.

### Analysis and modelling of sinale-cell allelic outcomes

To quantify cell-to-cell variability of allelic state in single tissue cells, we extracted colocalization data from our image analysis pipeline, and used this for further analysis. Using this data, we first compiled a quality control table for each experiment (Table S5) and excluded those where colocalization rates were <40% (4 out of 35 experiments). For all remaining data we combined replicates from the same genotype, and in the case of heterozygous data, combined results from C57BL/6J x JF1/Ms and JF1/Ms x C57BL/6J tissues. We then processed and visualised single-cell results using custom scripts written with R packages dplyr and ggplot2.

To determine how the observed data compared to random monoallelic expression (all-or-nothing scenario) or binomially distributed (coin-flip scenario) allelic calls we simulated those scenarios through modelling. For the binomial distribution we considered a null model wherein all heterozygous cells share the same allelic ratio, which was determined to be the overall allelic ratio observed at the population level. Then, for an experiment with overall estimated C57BL/6J allelic ratio equal to *p_BL6_* (above), we let 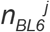 be the number of transcripts with C57BL/6J identity detected in cell *j* and 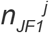 be the number the number transcripts with JF1/Ms identity detected in cell *j*. Under the null model, 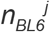 was drawn from a binomial with 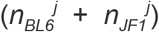 draws and probability *p_BL6_*. We simulated single-cell label counts for cells by drawing from these conditional null distributions for each cell 10,000 times. We then compared the negative log-likelihood of the observed data with the distribution of negative log-likelihoods of each simulation iteration.

To simulate random monoallelic expression each cell was assigned either a BL6 or a JF1 identity, based either on the majority of RNAs in a cell, or based on random assignment if both alleles had the same count. We then designated all RNAs in a given cell to the same allelic identity (eg a cell that originally contained 4 BL6 RNAs and 2 JF1 RNAs would be assigned a BL6 identity with 6 BL6 RNAs). Next, we randomly added “technical noise” 10,000 times, by changing some RNAs in the population to the opposite identity, based on the false positive rates measured in the original BL6 and JF1 homozygous populations. These steps were performed while keeping the final overall RNA assignments in the population the same as the original heterozygous population. For the 10,000 simulations we then calculated negative log likelihoods similarly as we did for the binomial distributions, assuming two separate null models for the BL6 and JF1 populations, whose parameters were determined by the original homozygous population.

To assess if the measured mRNA distributions were compatible with a transcriptional bursting scenario we used the negative binomial distribution to simulate expected mRNA counts (Raj et al. 2006). First, we determined the burst size and frequency of the BL6 and JF1 alleles separately, by using the moments method to determine r and p parameters of the negative binomial distribution based on the mean and variance of our measurements (where *p=mean/variance* and *r=mean*^*2/(variance-mean)*), from which we obtained the burst size and frequency using: *burst_size=(1-p)/p* and *burst_frequency=r*. We then generated 10,000 RNA counts for the two alleles separately by drawing from a distribution with the r and p parameters we had calculated. We visualized the obtained mRNA counts for both alleles individually using a randomly selected simulation, and also calculated the negative log-likelihood distribution of the 10,000 simulated datasets. Next, we randomly paired the data for the two alleles to generate “cells” with RNA counts from both alleles and calculated the correlation between the BL6 and JF1 counts in each of these modeled cells. In addition to using the negative binomial parameters that we had calculated from our data, we also tested a series of additional burst sizes (from 0.5 to 5 RNAs per burst) and repeated the entire analysis, which showed that our findings were consistent across a range of burst values (see Figure S6). Finally, to generate a model which included BL6-JF1 correlations that arise due to false assignments, we used the randomly paired data and changed some RNAs in the population to the opposite identity based on the false positive rates measured in the original BL6 and JF1 homozygous populations (one round of reassignments for each simulation). Following reassignment we again calculated the correlation between the BL6 and JF1 counts.

### Reproducible analyses

Raw and processed data, as well as scripts for all analyses presented in this paper, including all data extraction, processing, and graphing steps are freely accessible at the following url: https://www.dropbox.com/sh/5z8zsqdm48475zg/AACBwwW8UcdEnyx_T0968oOBa?dl=0 Our image analysis software (changeset: d278b7d0012282ecb318fde3bebbe3beaba62032) is available here:https://bitbucket.org/arjunrajlaboratory/rajlabimagetools/wiki/Home

## Figure Legends

**Figure S1.**
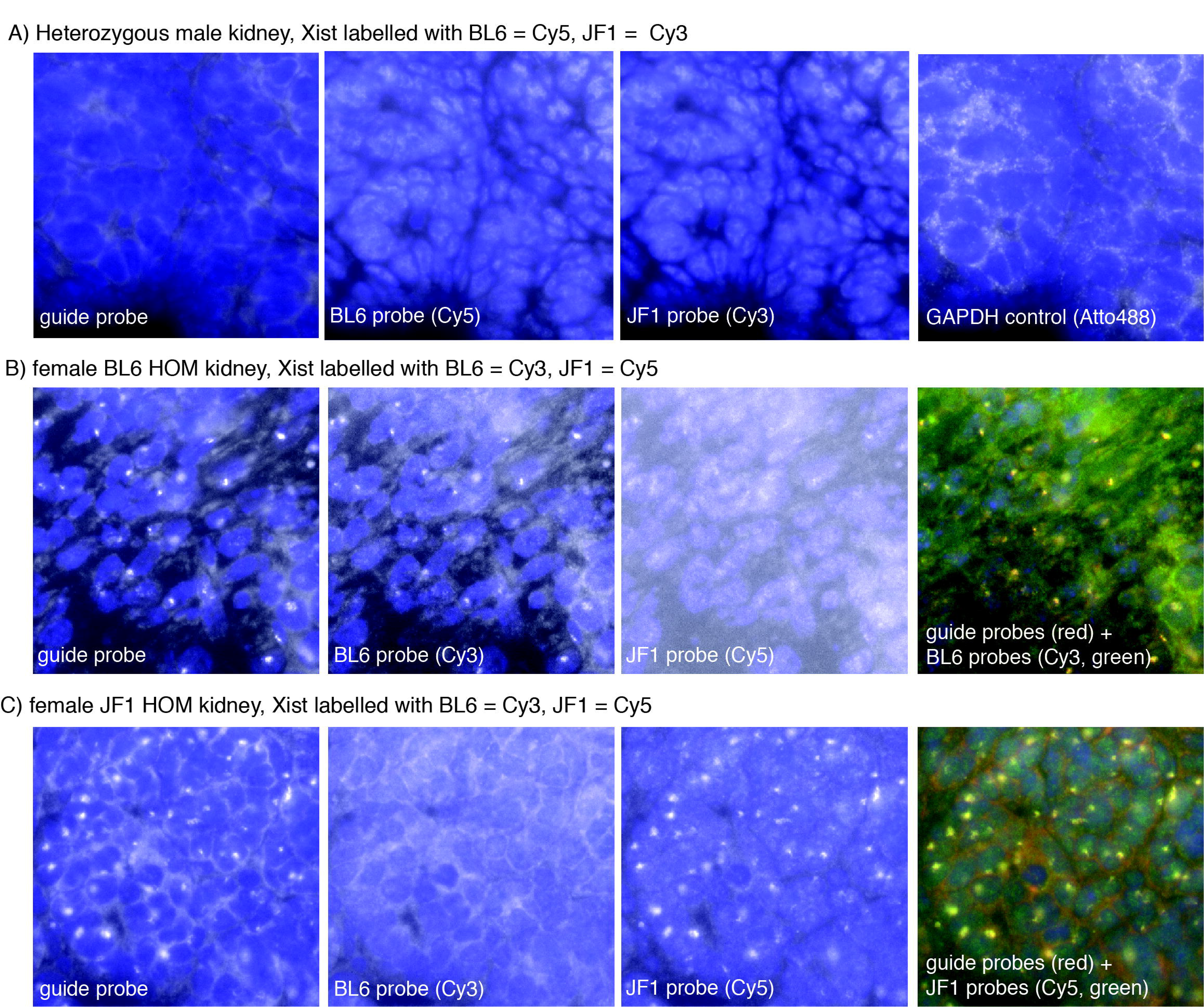
Xist RNA FISH signal in male and homozygous controls. **A.** Male heterozygous (BL6 x JF1) tissue was stained with probes for Xist (guide probe labelled with Cal fluor 610 (far left), BL6-specific probes labelled with Cy5 (middle left), JF1-specific probes with Cy3 (middle right)), none of which showed signal above background. Atto 488-labelled Gapdh probes were also included to verify absence of RNA degradation (far right). **B, C.** Female BL6 homozygous (B) and JF1 homozygous (C) tissues were stained with probes for Xist (guide probe labelled with Cal fluor610 (far left), BL6-specific probes labelled with Cy3 (middle left), JF1-specific probes with Cy5 (middle right)). Right-most image shows colocalization between guide foci and allele-specific foci as indicated.

**Figure S2.**
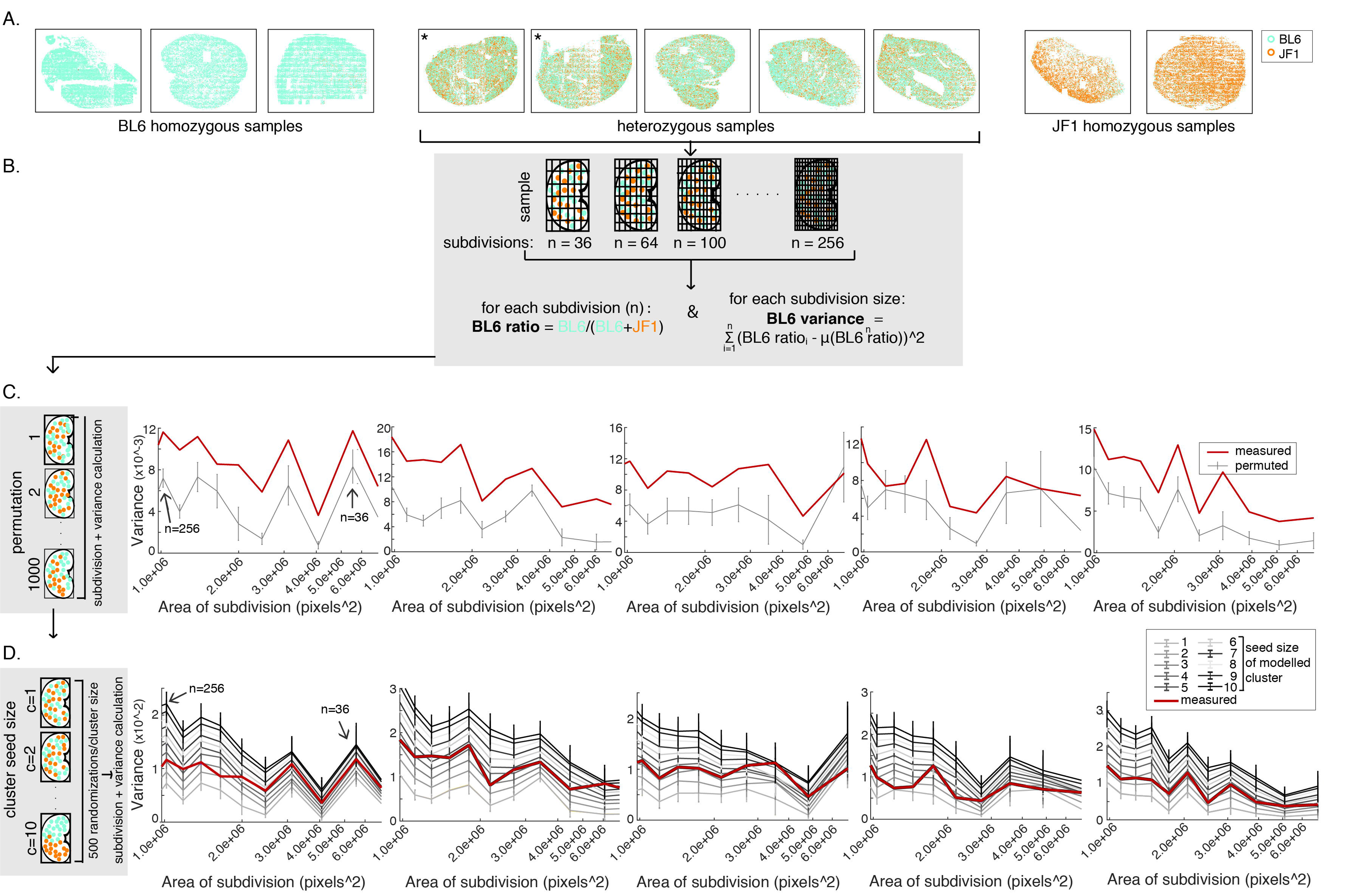
Allelic calls across whole tissue sections and modelling of spatial heterogeneity of Xist. **A.** Allelic assignments across different BL6 and JF1 homozygous (far left and far right) as well as heterozygous (middle) kidney sections. Two of the heterozygous kidney sections are technical replicates (different kidney sections from the same animal), which is indicated by an asterisk (“*”) . BL6 allelic assignment is depicted in turquoise, JF1 allelic assignment is depicted in orange. **B.** For all heterozygous samples we calculated spatial heterogeneity using a variance metric, the method of which is schematized: sections were subdivided into a grid, using increasingly smaller squares (from 8×8 to 16×16) and for each subdivision we calculated the ratio of BL6 Xist foci. For each grid we then also calculated the variance of the BL6 ratio across all squares of that grid. **C.** The measured variance (red line) was compared to the variances obtained for samples where we randomly permuted allelic assignments 1000 times (black line, error bars representing standard deviation of the modeled results). The graphs show the variance for subdivisions of different sizes, with both the area of the subdivisions and the size of the grid indicated. **D.** Measured variance (red line) was also compared to the variances of samples where we randomly placed different sized clusters (seeds) of allelically identical Xist foci in the tissue (lines in different shades of grey, error bars representing standard deviation of the modeled results). For each seed size we generated 500 randomizations, keeping the allelic ratio constant. For all heterozygous data shown in A, C and D the order of the samples is kept identical.

**Figure S3.**
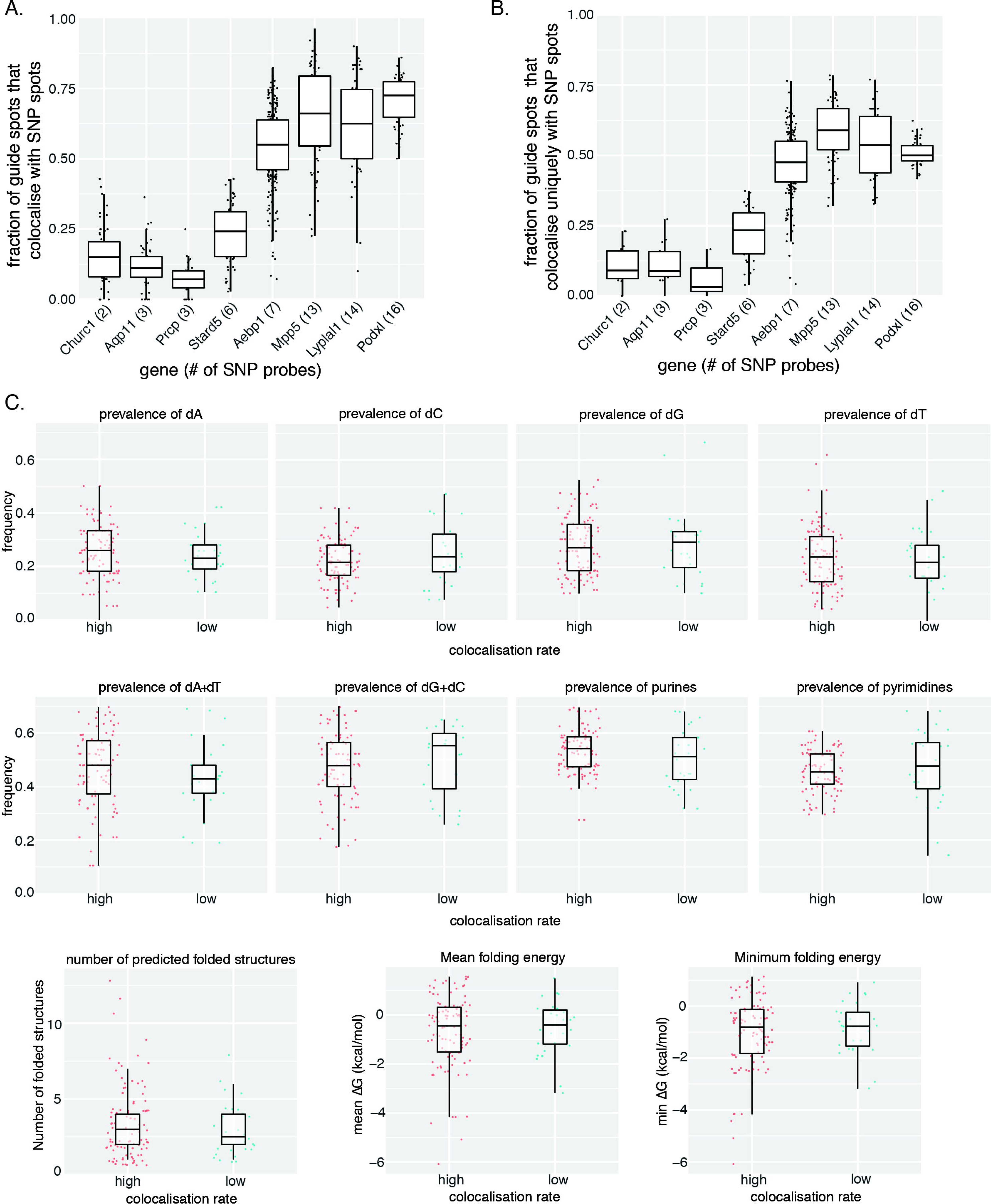
Colocalization rates and probe properties for autosomal allele-specific probes. **A, B.** Overall (A) and allele-specific (B) colocalization rates for different autosomal genes. Overall colocalization rates consider all guide spots that colocalize with either BL6 and/or JF1/C7 allele-specific signal, while allele-specific colocalization counts only those guide spots that colocalize uniquely with either BL6 or JF1/C7 probes. Each spot represent the colocalization rate in one area tested (typically 10-50 cells). All genes were detected with guide probes labelled with Cal fluor610, and the following allele-specific probes: *Aebp1, Mpp5* and *Podxl* BL6-specific probes labelled with Cy3, JF1-specific probes labelled with Cy5; *Churc1* and *Lyplal1* BL6-specific probes labelled with Cy5, JF1-specific probes labelled with Cy3; *Aqp11* and *Stard5* BL6-specific probes labelled with Cy3, probes for the C7 allele labelled with Cy5; *Prcp* BL6-specific probes labelled with Cy5, probes for the C7 allele labelled with Cy3. Genes are listed in increasing order of number of SNV probes utilized, which is indicated for each gene. **C.** Probe properties for probe sets with high (>50%) and low (<50%) mean overall colocalization rate. We compared prevalence of individual nucleotides (dA, dC, dG, dT - top row), nucleotides forming three hydrogen bonds (dC+dG) or two hydrogen bonds (dA+dT), purines (dA+dG) and pyrimidines (dC+dT) (middle row), as well as the number of folded structures predicted for each probe, mean and minimum folding energy for each probe (bottom row). For all plots, each spot represents the value obtained for a single probe.

**Figure S4.**
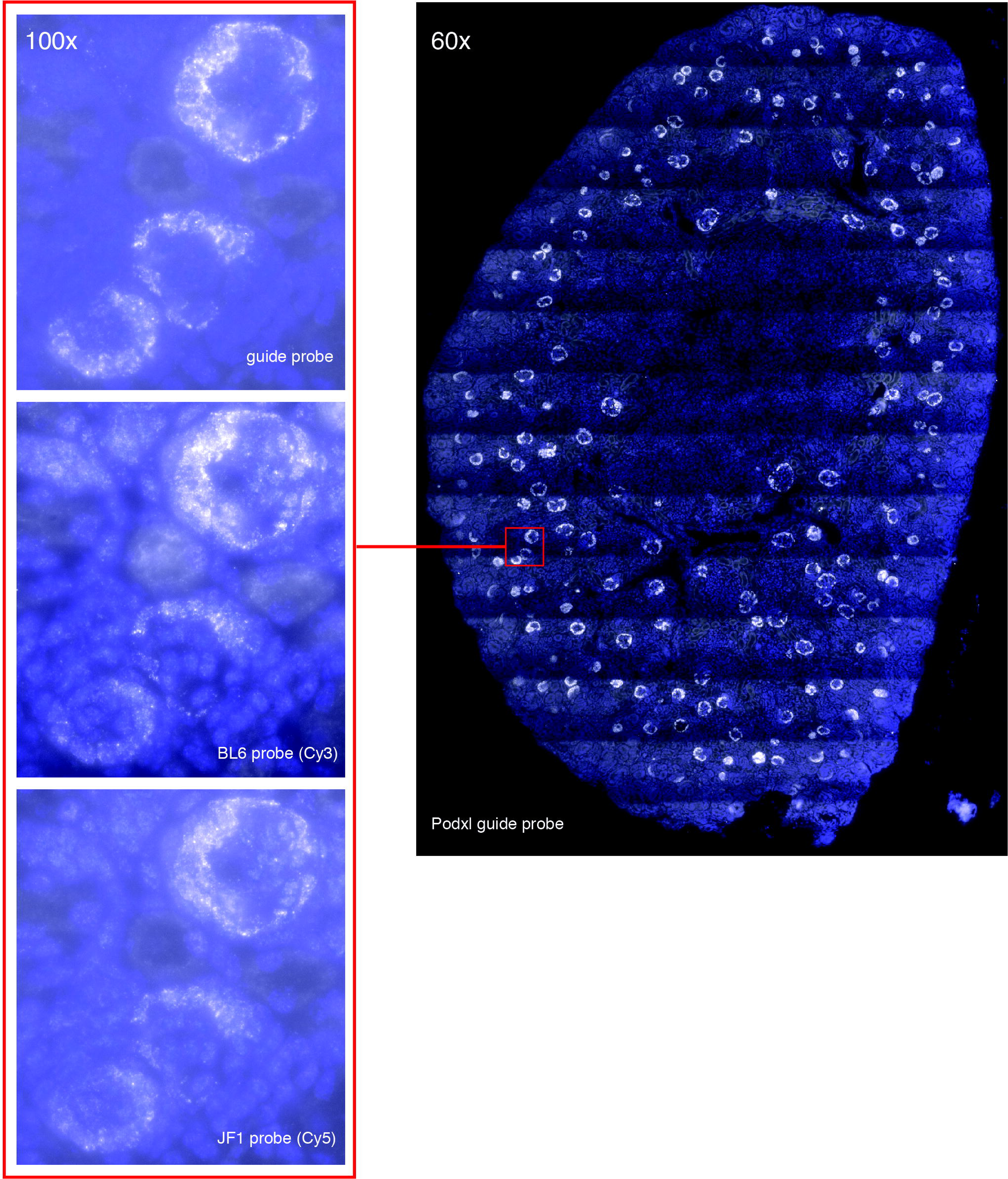
Allele-specific Podxl mRNA detection in mouse kidney sections. A tissue scan of the Podxl guide probe (labelled with Cal fluor 610) shows staining primarily in glomeruli, due to expression of the gene in podocytes (right). At 100x resolution these areas of high expression can clearly be distinguished from background when detecting fluorescence from the guide probe (left, top), as well as for the BL6 (left, middle) and JF1-specific probes (left, bottom). However, the high expression levels of the gene precludes the precise thresholding and detection of individual mRNA spots.

**Figure S5.**
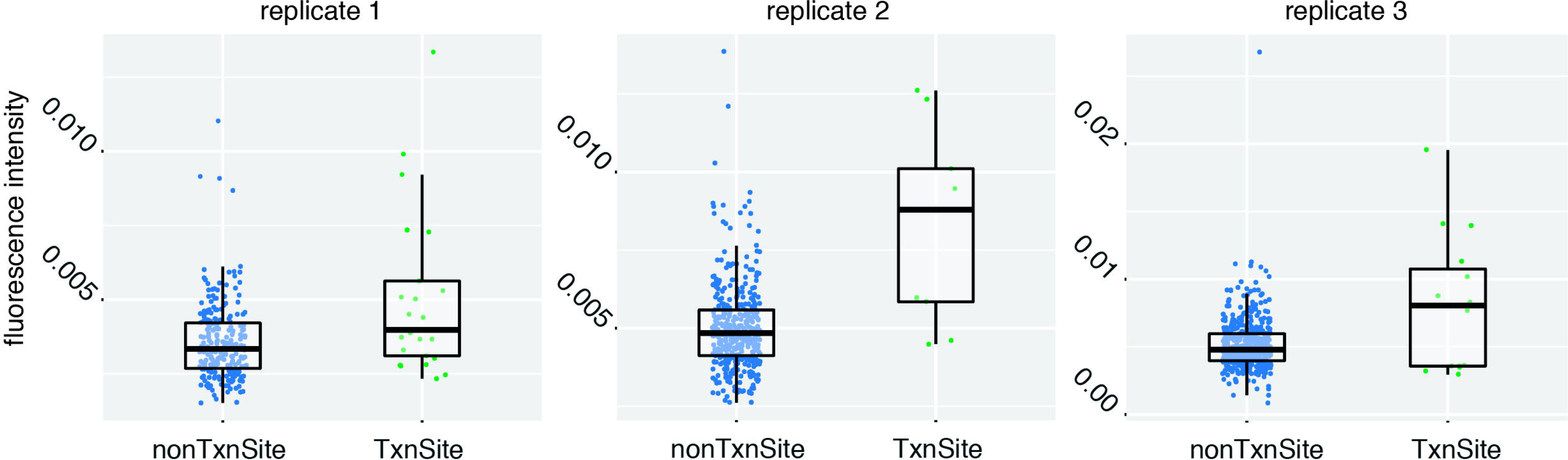
Fluorescence intensity of *Aebp1* mRNA at transcription sites and non-transcription sites. Each spot represents the intensity of a single mRNA guide spot (labelled with Cal fluor 610). Transcription sites were identified by overlap with intron probes labelled with Atto488 (replicate 1) or Alexa700 (replicate 2 and 3).

**Figure S6.**
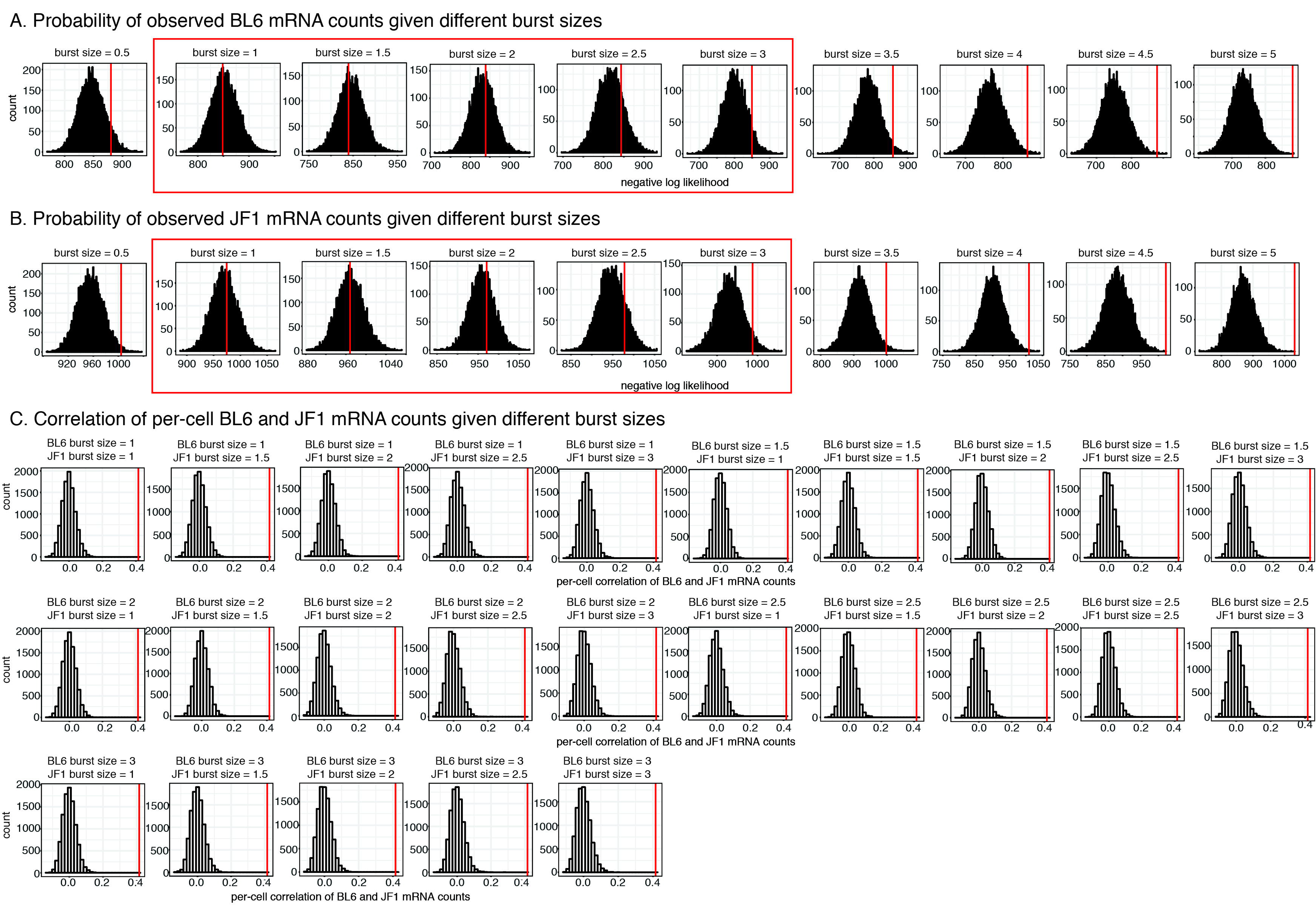
Probability and correlation of *Aebp1* BL6 and JF1 mRNA counts for simulations of transcriptional bursting using different burst sizes. **A, B.** Probability of observed BL6 (A) and JF1 (B) mRNA counts given different burst sizes. Burst sizes boxed in red showed a good fit for the two alleles individually and were used for correlation analysis. **C.** Correlation of per-cell BL6 and JF1 mRNA counts given the burst sizes that showed a good fit (boxed in red in A and B). Burst sizes of the alleles are indicated, and for each correlation analysis the simulations for BL6 and JF1 mRNA counts were paired up randomly. In all figures bar plots represent simulations, red line represents real data.

**Figure S7.**
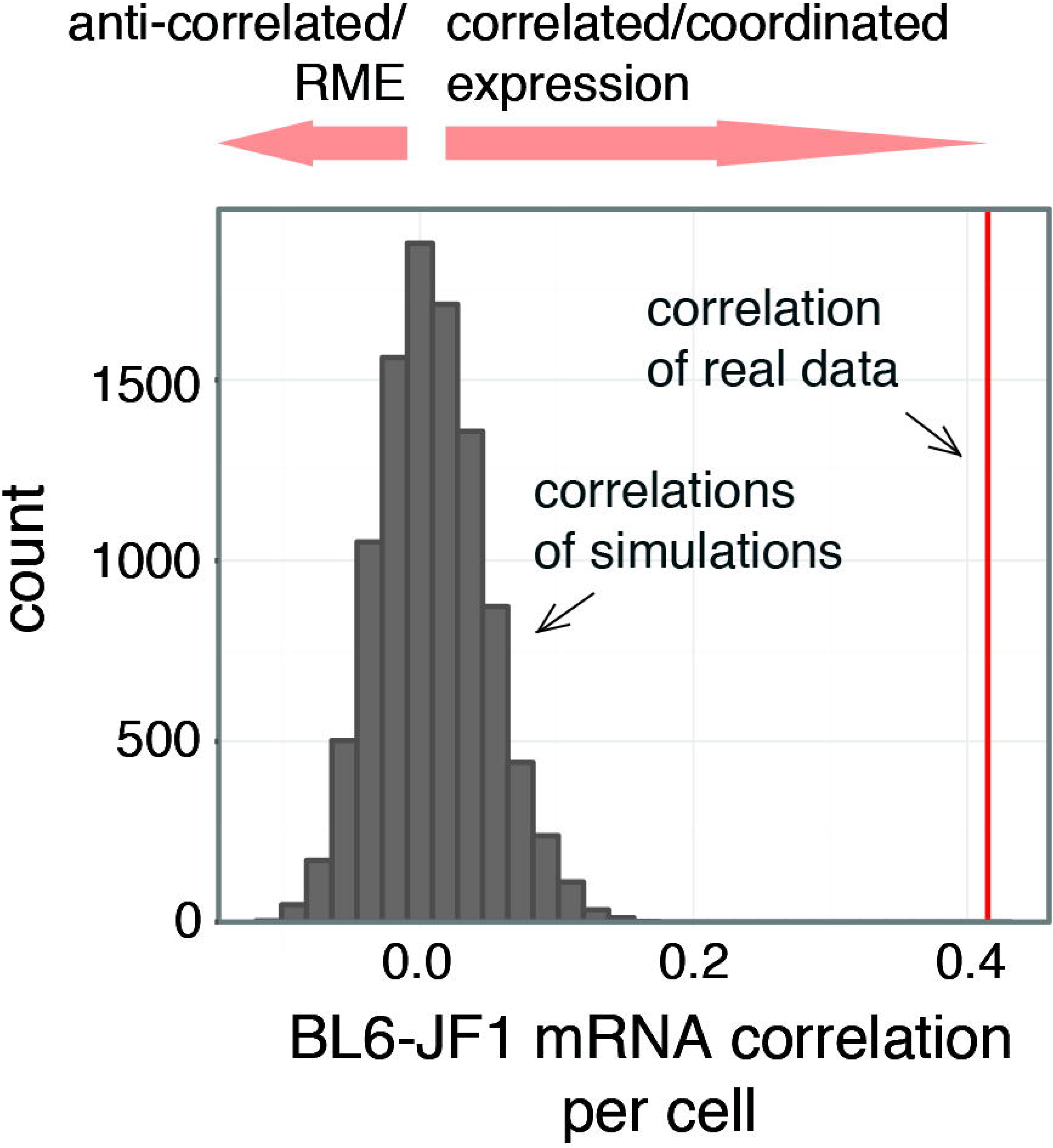
Correlation between BL6 and JF1 RNA counts in single cells when including false assignments in the simulations of bursty transcription. Correlation values for simulated counts in the case independent bursting from the two *Aebp1* alleles is shown as grey bar plot, correlation for real data is indicated as red line.

## Acknowledgements

We thank members of the Raj and Bartolomei laboratories for helpful suggestions and discussions. We particularly thank Chris Krapp for assistance with animal care and husbandry, Joanne Thorvaldsen for suggestions about Xist analysis, Paul Ginart for his advice on methodology and Rohit Gupte for help with transcription site code. A.R. and M.S.B. acknowledge support from from NIH/NIBIB R33 EB019767, A.R. from NSF CAREER Award 1350601, NIH 4DN U01 HL129998 and NIH Center for Photogenomics (RM1 HG007743). O.S. received funding from EMBO long-term fellowship ALTF 691–2014 and Human Frontier Science Program fellowship LT000919/2016-L. I.A.M. acknowledges funding from NIH/NIGMS T32GM007170 (University of Pennsylvania MSTP), NIH/NHGRI T32HG000046, and NIH/NINDS F30NS100595. J.M.K. acknowledges funding from NIH K08 CA193915, St. Baldrick’s Foundation, and Margaret Q. Landenberger Foundation. Sectioning was performed at the Penn Center for Musculoskeletal Disorders Histology Core (NIH P30-AR06919).

## Author contributions

O.S., M.S.B. and A.R. conceived the project, O.S. and J.M.K. performed experiments, O.S., M.C., I.A.M. analyzed the data, O.S. and A.R. wrote the paper.

